# U.S. Visa Bureaucracy and Its Burdens Among Early Career Scholars

**DOI:** 10.1101/2025.07.28.667220

**Authors:** Michel Nofal, Raquel Ferrer-Espada, Payel Ganguly, Mayank Chugh

**Affiliations:** Harvard Medical Postdoc Association, 25 Shattuck Street, Boston, 02115; Wyss Institute at Harvard, 201 Brookline Avenue, Boston, MA 02215; Department of Systems Biology, Harvard Medical School, Boston, MA 02115; Department of Cell Biology, Harvard Medical School, Boston, MA 02115; ReForm Lab, Biology Department, College of William and Mary, Williamsburg, VA, 23185; Gender, Sexuality, and Women’s Studies Program, College of William and Mary, Williamsburg, VA, 23185

## Abstract

Foreign-born research scholars on temporary visas are essential to the U.S. scientific workforce and economy, yet they face systemic challenges shaped by restrictive immigration policies. While scholars across disciplines have long analyzed the evolution of U.S. immigration regimes and citizenship privilege, there remains a critical lack of quantitative data on the financial, temporal, and psychological burdens associated with visa bureaucracy. Here we present original survey data from more than 700 international postdoctoral researchers at Harvard Medical School (HMS) and its affiliated institutions, revealing how visa-related burdens directly impact scholars and their research. For example, over 40% of scholars reported spending more than a month in their home countries to renew U.S. visas, with 5% spending more than six months. Scholars from Asia experienced longer delays and higher costs than those from Europe. Seventy-five percent of respondents reported mental health challenges related to visa stress, often feeling trapped at work, with some requiring medication and time off. Our findings underscore the urgent need for targeted institutional support to promote research productivity, equitable opportunity, and mental well-being. As the U.S. repositions itself as a global scientific leader, such reforms will be critical to recruiting and retaining top talent and advancing equity in science.

## Introduction

International students, scholars, and workers play an essential role in the U.S. STEMM (Science, Technology, Engineering, Mathematics, and Medicine) ecosystem. Nearly 40% of U.S. doctoral graduates and 55% of postdoctoral researchers are on temporary visas (*Bernstein 2022, Agarwal 2023, NSB 2022*), and immigrants have founded over half of U.S. tech companies valued at $1 billion or more (*Keith 2021*). Immigrant scientists have received 40% of U.S. Nobel Prizes in STEMM since 2000 and comprise more than half of full-time Science & Engineering (S&E) faculty (*NFAP 2023*). Their contributions generate an estimated $460B in labor value and added $40B to the U.S. economy in 2022–2023 alone (*Fwd 2024*).

Most foreign-born researchers enter the U.S. through temporary visas such as F-1 (students), J-1 (exchange scholars or short-term trainees), or O-1 (exceptional ability), later transitioning to H-1B status for employment. Created in 1990, the annual statutory cap for H1B visas is 65,000 with an additional 20,000 visas for graduates who hold advanced degrees, such as master’s or doctorate, from a U.S. higher education institution (*AIC 2025, Peri 2020*). In academia, scholars employed by non-profit higher education institutions are exempt from the statutory cap allowing more flexibility in hiring talented researchers and educators. Still, this pathway remains precarious, bureaucratically complex, and highly unequal across global regions.

Visas and citizenship have long been an area of academic inquiry. Citizenship privilege is inextricably linked to occupational mobility and migration of people (*Tan 2024, Chugh 2024, Castles 2004, Glick 2013, Mau 2015*). In academia, mobility is a crucial attribute for academic scholarship and practice to build a successful and competitive career. Numerous studies have provided data, evidence, and theories for social determinants and implications of international mobility within academia (*Fernandez-Zubieta 2013, Van Noorden 2012, Jonkers 2013, Scellato 2017*). Despite that, comprehensive data on visas and citizenship of mobile scholars remains insufficient. For instance, while National Science Foundation (NSF) annual surveys mention the proportion of doctoral graduates and postdoctoral scholars on temporary visas in the US, they do not mention their visa type and citizenship (*NCSES 2024*). The U.S. Citizenship and Immigration services (USCIS) does reveal H1B visa holder citizenship data; however, these datasets likely underrepresent foreign-born STEMM scholars in academia.

Despite their significant contributions to the STEMM enterprise and U.S. economy, immigrant researchers often face suboptimal treatment. Numerous reports show mistreatment, harassment, and exploitation of foreign-born scholars, especially among early career researchers (ECRs) like undergraduate and graduate students and postdoctoral scholars on temporary visas (*Hafid 2022, Wadman 2017, Kahn 2024, Hayter 2019, Woolston 2020*). A recent report demonstrated that ECRs on temporary visas are more productive but receive lesser mentorship, monetary compensation, and career support than U.S. citizens and permanent residents. Findings that are consistent with previous reports by Sigma Xi and National Academies of Sciences (NAS) (*Kahn 2024, Davis 2009, NRC 2005*). In addition to workplace treatment, immigration policies and visa bureaucracy exacerbate the global north-south knowledge divide, thus, reinforcing patterns of citizenship privilege (*Chugh 2024, Arias Cubas 2024, Owusu-Gyamfi 2024*).

Although visa-related challenges, such as renewal and processing delays, are often assumed to impact all temporary holders, scholars from the global south face disproportionately greater hardships in visa acquisition and taxation on their personal and professional lives (*Chugh 2024, Owusu-Gyamfi 2024, Nshemereirwe 2018, Normile 2021*). For example, a recent report shows that the denial rate for F-1 visas is approximately 54% for African students, followed by 36% for Asian students compared to 9% for European students (*Bhandari 2023*). Another study by Chatterjee et al. analyzed roughly 10,000 biomedical doctoral and postdoctoral scholars in U.S. academic institutions with an intersectionality approach and found that non-citizen women exhibit less career self-efficacy (*Chatterjee 2024*).

In this study, we present data on the impact of U.S. visa bureaucracy on approximately 750 international postdocs at Harvard Medical School (HMS) and its affiliated institutions. Contrary to common assumptions, this work offers the first quantitative analysis of visa types, renewal timelines and costs. We also show that visa-related administrative burdens not only exacerbate psychological stress but also restrict long-term career planning, stability, and professional advancement. Our findings underscore the need for more equitable institutional policies and challenge the notion of international scholars as a monolithic group.

## Materials and Methods

In this study, we determined the geographical composition of the postdoctoral workforce at Harvard Medical School (HMS) and its affiliated institutions with respect to citizenship and visa bureaucracy. Second, we identified the administrative burden of U.S. visa bureaucracy on postdoctoral researchers’ mental wellbeing, career outcomes, and support they receive from employing labs or institutions. We focused on the postdoctoral workforce for several reasons. a) The postdoctoral workforce is the primary backbone of research-intensive institutions and represents the highest proportion of international scholars within the US higher education system. b) Postdocs are among the most vulnerable academic populations, facing challenges such as low job security, limited compensation, insufficient institutional support, and often difficult workplace conditions. c) There is a lack of institutional transparency and data about postdocs likely because of temporary position status and several differing job titles. (d) The HMS postdoc population is among the largest in the country, thus, serving as a model subset to study the trends.

### Data collection

A survey was developed using web-based software Qualtrics. The Harvard Longwood Campus IRB determined this study to be exempt from IRB review (Exempt Study; Protocol #IRB23-1379). The survey contained 51 questions. The survey began with a consent form, and participation in each question was entirely voluntary. To ensure anonymity, respondents’ IP addresses, location data, and contact information were not recorded.

The anonymous survey was disseminated via email by the Harvard Medical Postdoc Association (HMPA), Harvard Medical School (HMS), Harvard School of Dental Health (HSDH), Office for Postdoctoral Fellows (OPF), Harvard Catalyst, various school departments, local postdoc associations, and through flyers placed at the target institutions. The constituents of the survey were only postdocs employed at Harvard Medical School (HMS) and its affiliated institutions. HMS-affiliated institutions include, Beth Israel Deaconess Medical Center (BIDMC), Boston Children’s Hospital (BCH), Brigham and Women’s Hospital (BWH), Massachusetts Eye and Ear with Schepens Eye Research Institute (MEE), Massachusetts General Hospital (MGH), McLean Hospital (MH), Dana-Farber Cancer Institute (DFCI), Harvard Pilgrim Health Care Institute (HPHCI), Joslin Diabetes Center, Judge Baker Children’s Center, Cambridge Health Alliance (CHA), Mount Auburn Hospital, Hebrew Senior Life, Spaulding Rehabilitation Hospital (SPH) and VA Boston Healthcare System. The data was collected from January 09, 2024 till March 22, 2024. As one of the drawbacks of working with postdoctoral researchers, systematic institutional data was not available.

### Analysis

Survey responses were extracted from Qualtrics as.csv files. A custom Python script was used to clean the data by removing incomplete submissions and formatting inconsistencies. Data visualization was performed using custom written python scripts and null data points were excluded. All figures were created in Adobe Illustrator 2025.

## Results

### Geographic and Visa Demographics of HMS Postdoctoral Scholars

A total of 735 foreign-born postdocs at HMS and its affiliated institutions answered the survey (**Figure 1a-b**). The majority—67.44%—were J1 visa holders, typically associated with research or educational exchange programs (*For visa type details, see Appendix A*). 14.9% held H1B visas, commonly granted for specialized roles in fields such as technology and research. Additionally, 8.6% were on F1/OPT visas generally used by international students or recent graduates seeking temporary work experience (**Figure 1c**). A further 2.0% fell under other visa categories, while 6.7% were green card holders, reflecting permanent resident status in the U.S. A small proportion, 0.3%, held O1 visas, designated for individuals with extraordinary ability.

**Figure 1:**
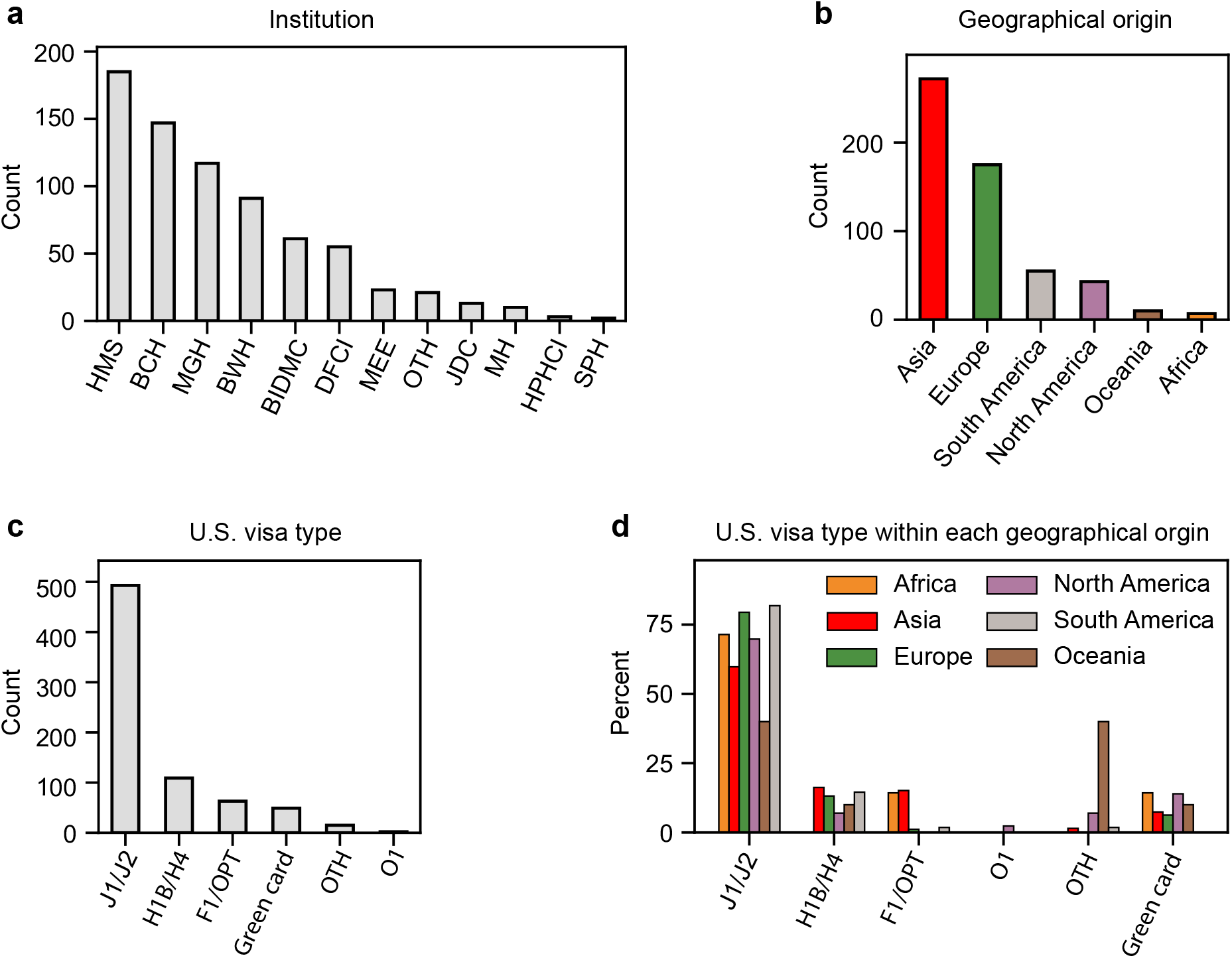
Institution and U.S. visa type demographics among HMS postdocs. a) Number of foreign-born postdocs on temporary U.S. visas who completed the survey at each participating institution. HMS, Harvard Medical School; BCH, Boston Children’s Hospital; MGH, Massachusetts General Hospital; BIDMC, Beth Israel Deaconess Medical Center; DFCI, Dana-Farber Cancer Institute; MEE, Massachusetts Eye and Ear; JDC, Joslin Diabetes Center; MH, McLean Hospital; HPHCI, Harvard Pilgrim Health Care Institute; SPH, Spaulding Rehabilitation Hospital; OTH, Other (comprising a few smaller institutions listed in the Methods section). b) Number of postdocs based on their geographical origin. c) Number of postdocs by U.S. visa type. d) Percentage of postdocs by U.S. visa type within each geographical origin, a proxy for citizenship.

Respondents’ country of origin was categorized into regions based on the classifications provided by the Office of Homeland Security Statistics (*OHSS, see references*). Of all participants, 56.9% voluntarily shared their demographic information. Among them 48.6% were from Asia, 30.9% from Europe, 9.8% from South America, 7.7% from North America, 1.8% from Oceania, and 1.3% from Africa (**Figure 1c-d**). When considering temporary non-immigrant U.S. residents only on J1 exchange visitor visa (majority of respondents), our reported trend is almost consistent with OHSS (43% Asia, 33% Europe, 10% North America, 9% South America, 6% Other).

### Financial and Temporal Burdens of Visas Are Deeply Unequal and Disruptive

We asked the international postdocs about the financial and logistical challenges involved in obtaining and renewing their visas. Across all visa types and regions of origin, nearly 30% of respondents reported renewing their visas annually, while 20% renewed every two years. The remaining respondents renewed less frequently. Strikingly, over 40% of respondents reported spending more than a month on the visa renewal process (**Figure 2a-b**). This timeline includes completing the application, gathering required documentation (such as translated materials), and traveling to a U.S. embassy or consulate—often in their home countries, as U.S. visas must be renewed from abroad. Even among those able to complete the process in under a month, the average cost was around $1,000. For 22% of respondents, each renewal cost over $2,000, with some reporting expenses exceeding $5,000 (**Figure 2c-d**). Due to the limited number of respondents from Africa, Oceania, and North and South America, in certain categories and to protect respondent anonymity we grouped these regions together under the category ‘Other’. On average, postdocs with Asian citizenship reported longer visa processing times and higher renewal costs compared to their peers from Europe and other regions.

**Figure 2:**
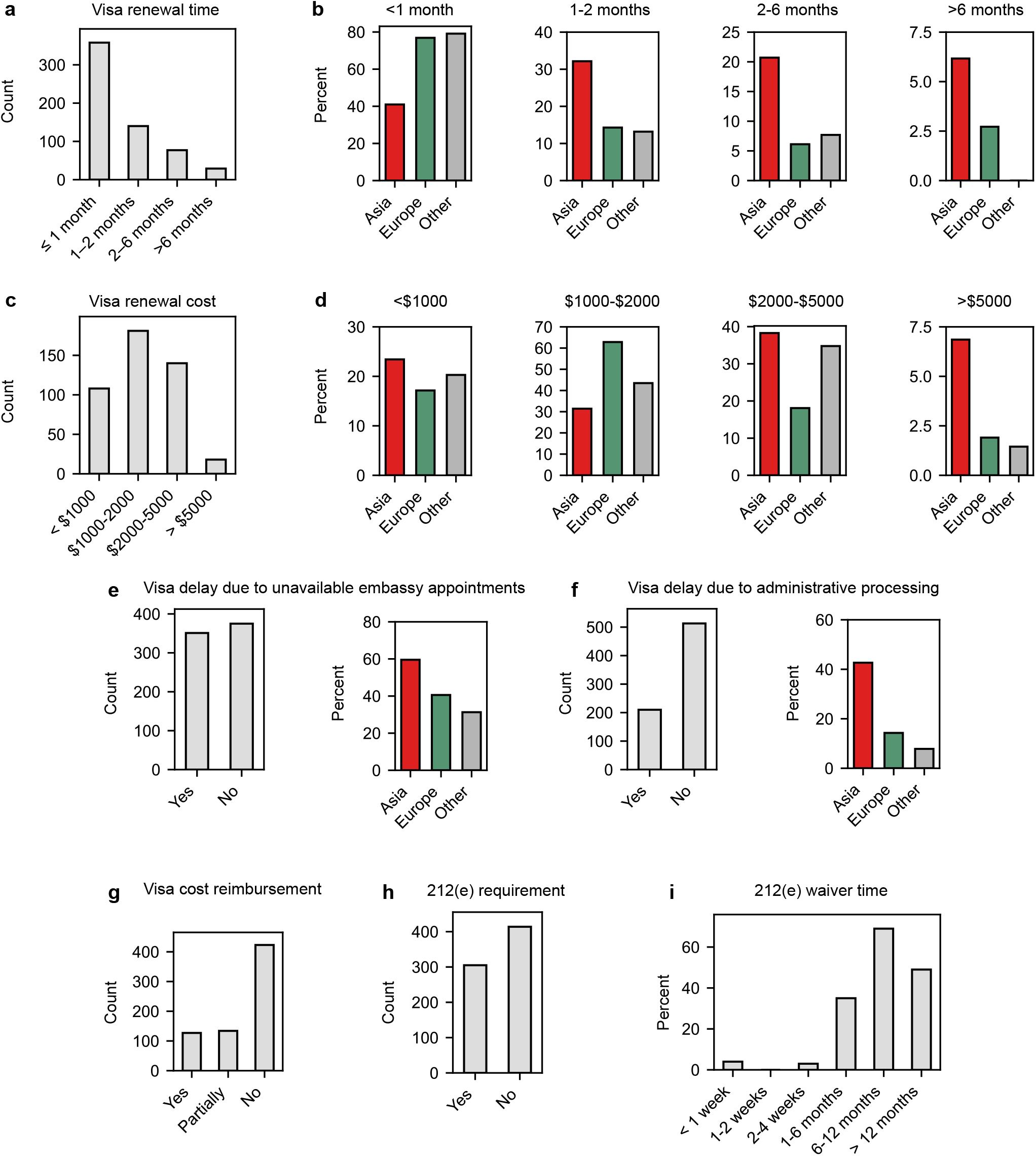
Financial and logistical burdens of visa bureaucracy on HMS postdocs holding temporary U.S. visas with a skew towards Asian citizenship. a) Visa renewal times by total number of postdocs. b) Visa renewal times by postdoc percentage within geographical origin. The other bar includes responses from North America, South America, Oceania, and Africa, due to significantly limited data. c) Visa costs borne directly by postdocs. d) Visa costs within geographical origin of postdocs, shown as percent postdoc population. e) Visa delays experienced due to unavailable embassy appointments at U.S. embassies and geographical origin. f) Visa delays administrative processing and geographical origin. g) Proportion of postdocs reimbursed for visa costs by host institutions. h) Number of postdocs subject to Section 212(e) under U.S. immigration law. i) Time spent by postdocs in obtaining a 212(e) waiver from their home countries.

Nearly 50% of all respondents delayed or altered travel plans due to embassy appointment unavailability (**Figure 2e-f**); the figure rose to around 60% among Asian postdocs. Nearly 29% reported visa delays exceeding 10 days, and 2.6% experienced outright denials. Of those affected, only 21.4% received official explanations. Importantly, 62% of respondents did not get any reimbursement from their institutions and only 32% received partial reimbursement on their out-of-pocket visa costs (**Figure 2g**).

Visa-related delays are not minor inconveniences—they are avoidable, structural disruptions with far-reaching consequences for scholars, academic institutions, and the broader U.S. research enterprise (*The Risk of Exclusion, 2003; Woolston, 2017; Morello, 2017; Udesky, 2025*). For ECRs, these delays can halt time-sensitive experiments, derail grant submissions, interrupt teaching and mentoring commitments, and block access to educational opportunities they have already paid for. They also limit participation in essential professional development, including conference presentations and collaborative research, activities critical for academic visibility and advancement. For faculty and institutions, these delays can result in inefficient use of public research funds and contribute to the broader loss of highly skilled talent essential to scientific advancement and economic growth.

A substantial 42.4% of postdocs in our sample were subject to the two-year home residency requirement under Section 212(e) of U.S. immigration law **(Figure 2h)**. This provision mandates that scholars on J-1 visas must return to their home country and physically reside there for two years following the completion of their J-1 status before becoming eligible to apply for H-1B, L, or other immigrant visa categories. Undoubtedly, a two-year interruption poses a major disruption to scientific research and can significantly derail a scholar’s career trajectory. Since the maximum duration of a J-1 visa is five years, many postdocs seek a waiver of the 212(e) requirement from their home governments in order to continue their research careers in the U.S. Among those who applied for a waiver, 30% reported that the process took over one year (**Figure 2i**), with longer durations particularly prevalent among scholars from Asian countries. The financial burden of the waiver process also varied: 45.6% of applicants spent less than $500, 39.8% spent more than $500, and 14.5% reported incurring no costs.

### Psychological Toll Underlie Most Foreign-born Postdocs on U.S. Visa

Survey participants were also asked about the impact of their visa status on their career trajectory as well as mental health (**Figure 3**). Visa-related stress had significant mental health contribution on postdocs wellbeing. A striking 75% of respondents reported experiencing mental health challenges (defined in the survey as anxiety, insomnia, panic attacks, low motivation, mood swings, low appetite, and low self-worth) linked to the visa application process (**Figure 3a**). Only 3% reported no impact, and 22% preferred not to answer. Among individuals who acknowledge mental health challenges, 11% were prescribed medication and 29% took time off work (**Figure 3b-c**). Visa constraints also limited postdoc mobility from toxic work environments. Nearly half of respondents felt unable to leave due to visa restrictions. Encouragingly, while 32% were aware and trusted their institution’s mental health support services, 18% did not (**Figure 3d**). When asked about experiences of harassment or discrimination related to visa status, nearly 17% experienced it (**Figure 3e**), yet only 5% formally reported it.

**Figure 3:**
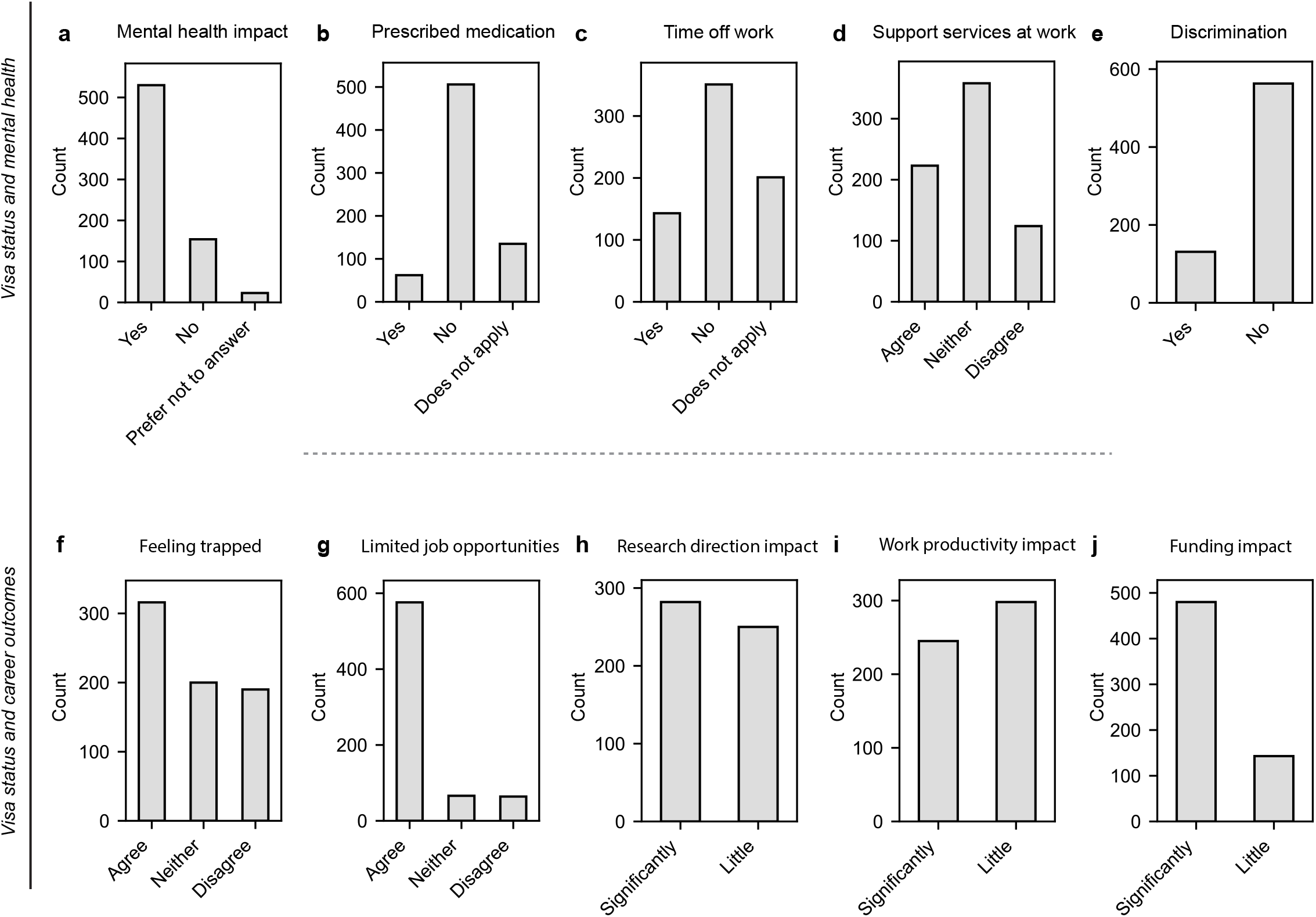
Psychological burdens of visa-holding HMS postdocs reveal significant impacts on mental health and career outcomes. a–e) Proportion of postdocs who: (a) experienced mental health challenges, (b) needed prescription medication, (c) took a break from work due to mental illness, (d) were informed about mental health services at their institutions, and (e) experienced discrimination in the workplace due to their visa status. f–j) Proportion of postdocs who reported visa-related barriers to: (f) advancing their careers, (g) pursuing job opportunities, (h) maintaining their original research direction, (i) sustaining scientific productivity, and (j) accessing research funding.

We found visa constraints also shaped postdocs career trajectories. A significant majority, 44.7% felt ‘*trapped*’ in their postdoctoral positions due to visa status, and 81.5% reported limited job opportunities (**Figure 3f-g**). Similar sentiment of academic entrapment due to visa has been reported from a large study of U.S. PhD graduates who could not work in technology start-ups (*Roach 2019*). Additionally, nearly 50% indicated visa status influenced research direction; 45.1% disclosed a negative effect on their scientific productivity (**Figure 3h-i**). Most notably, 80% reported that visa limitations restricted their access to funding (**Figure 3j**). These findings echo earlier surveys: a 2004 national report found 20% of nonresident postdocs curtailed travel due to reentry concerns, with over 20% encountering reentry problems and 2% facing serious issues (*NRC 2025*). We do note that these visas linked psychological burdens on HMS postdocs may not be exclusive and contributed by additional factors such as funding and workplace conditions.

These findings underscore the challenges faced by postdocs on temporary visas, including effect on research productivity, struggles with mental health, limited job mobility, and a reluctance to report workplace mistreatment, all tied to their temporary visa status, highlighting stigma and systemic vulnerabilities.

### Discussion and Conclusion

While scholars across disciplines have long documented the evolution of U.S. immigration regimes and immigrant experiences (*Goldin 1994; Joseph 2017; Cox 2020; Moynihan 2022*), there remains a paucity of quantitative evidence capturing how immigrants navigate bureaucratic systems. The 2023 Kaiser Family Foundation KFF/LA Times immigrant survey (*Schumacher 2023*) offers rare large-scale insight but lacks analysis of visa types, renewal times, and costs, which constitutes administrative burden—an omission that limits its utility in policy reform.

In this case study conducted at HMS and its affiliated institutions, we investigated the US visa bureaucracy and its burdens on international postdocs, an important and vulnerable academic population. Despite serving as essential contributors to U.S. scientific output, postdocs on temporary visas face systemic precarity—job insecurity, unstable funding, discrimination, and opaque career pathways (*Curtis 1969; Powell 2015; Woolston 2020*).

Efforts to improve national-level data collection by the National Postdoctoral Association (NPA) and NSF/NIH-sponsored surveys have helped track demographic trends and experiences, but critical gaps remain. For instance, National Center for Science Engineering and Statistics (NCSES) datasets distinguish between U.S. citizens and non-citizens but often omit finer-grained information on country of origin and visa type, despite its strong correlation with global mobility and access to opportunity (*Chugh 2024; Talavera-Soza 2023*). This missing granularity obscures the disproportionate burden borne by scholars from the global south. Existing studies already indicate that these scholars face higher visa denial rates and career self-efficacy challenges (*Chatterjee 2024; Bhandari 2023*).

Our findings are a first attempt in underscoring those disparities. Most HMS international postdocs spend a month and $1,000–$2,000 per visa renewal, with postdocs from Asia facing even higher burdens, up to 6 months and up to $5,000. These costs are rarely reimbursed by institutions. While some of this can be attributed to distance-related expenses, the more pressing concern is the inequity across countries. *Recchi et al. 2021* quantify this global visa cost divide, showing that citizens from Sub-Saharan Africa and South Asia must work significantly longer than Europeans to cover equivalent visa fees.

More importantly, these extended visa delays, up to 6 months, inevitably disrupt research continuity and timelines: missed grant deadlines, teaching, conference presentations, lab meetings, and time-sensitive experiments can set back entire projects. For scholars in lab-based disciplines, such disruptions can cause data loss, stalled projects, or failed collaborations. In the competitive academic job market, these cumulative disruptions may derail publication schedules, delay or disqualify for job applications, or sever collaborative opportunities—outcomes that can drastically alter a scholar’s career trajectory. Moreover, visa-related travel restrictions often force researchers to decline invitations to international meetings, diminishing their visibility and professional networks. Prior research has shown that such administrative barriers systematically hinder the productivity and advancement of foreign-born scientists (*Franzoni et al., 2012; Stephan, 2012*).

These financial and temporal burdens are compounded by psychological costs. Nearly 75% of postdocs reported mental health impacts, including anxiety and burnout, with 30% taking time off work. About half reported negative effects on research productivity and direction. Asian postdocs were particularly affected, though limited representation from other regions constrained further analysis. The cognitive costs could also be related to structural or organizational stressors, such as workplace conditions or access to health care benefits that are reportedly a longstanding concern among postdoc fellows (*Ferguson 2021*). These patterns mirror broader findings linking immigration-related stress to chronic health disparities (*Dunjacik 2023*).

Visa constraints also shape postdoctoral career mobility. Half of respondents felt ‘*trapped*’ in their roles, and 80% reported restricted job prospects due to visa status. U.S. immigration law legally binds nonimmigrant scholars to their host institution, often forcing them to endure harmful work environments to avoid leaving the country (*Nam 2023*). *Roach et al. 2019* finds that STEM PhDs on F-1 visas face barriers to entering start-ups, a challenge likely shared by J-1 visa holders, who form most of our dataset. The J-1 system, nominally for cultural exchange, has increasingly become a low-cost labor mechanism, with no annual caps, no employer payroll tax, less built-in accountability by the federal departments and thus, lacking the protections typical of other employment visas (*Flora 2020*). This vulnerability is intensified by the 212(e) two-year home residency requirement. Nearly half of respondents were subject to it, with Asian and African scholars most affected, often requiring a year or more to secure a waiver to the rule. The time and financial costs associated with this process further likely constrain mobility and career advancement.

It is important to recognize that the extended and often redundant U.S. visa bureaucracy not only affects countless ECRs but also serves as a barrier to the global circulation of talent and expertise, ultimately hindering scientific advancement and economic development. Although many scholars have shared their lived experiences navigating these challenges (*Morello 2017; Woolston 2017*), there remains a critical need for systematic analysis and policy engagement. This study represents a meaningful step toward addressing these structural barriers through empirical evidence.

In sum, our study quantitatively documents the administrative burdens faced by international postdocs on temporary visas at HMS and affiliated institutions, thus, demonstrating how citizenship status mediates access to opportunity, stability, and well-being in the U.S. research enterprise. These findings extend the administrative burden framework to the postdoctoral context and call for more equitable institutional policies and immigration reforms to support the international scientific workforce. Without targeted support and institutional flexibility hosting foreign-born scholars, these burdens risk entrenching global inequities in scientific training and leadership, effectively sidelining some of the most talented researchers from contributing fully to the U.S. research enterprise.

## Limitations

First, the sample is limited to HMS and affiliated institutions, constraining generalizability. Nonetheless, the observed trends align with qualitative accounts of immigrant postdocs and underscore systemic policy shortcomings (*Prentice 2016; Agbor 2021; Klotz 2023*). A national-level study, ideally in collaboration with the National Postdoctoral Association (NPA), is needed to validate and expand these findings. Second, the survey did not comprehensively capture variables such as mentoring, compensation, job titles, promotion pathways, postdoc duration, and career trajectories, each of which may be shaped by citizenship and thus, immigration status. In follow-up studies, we aim to expand our analysis by integrating national postdoctoral data to establish citizenship as a central variable in intersectional analyses of scholars’ recruitment, retention, and career outcomes. Such analyses are critical for two reasons: (1) cultural and geographical contexts shape expectations around gender roles and career progression, and (2) women disproportionately face work–family conflict and role spillover (*Webster 2021; Parker 2015*). Future research should therefore incorporate work–family dynamics to build a more nuanced understanding of how intersecting identities structure postdoctoral career outcomes and success.

## Acknowledgements

The authors are grateful to Jim Gould and Michaela Tally at the Office of Postdoctoral Fellows of the HMS for their unwavering support, funding and resources. The authors also thank members of the Harvard Medical Postdoc Association (HMPA) Governing Board 2022-2024. The authors are grateful to Tiffany Joseph at Northeastern University, Heidi Prozesky at Stellenbosch University, Crystal Peoples, Helen Murphy, and the ReForm Lab at the College of William & Mary for feedback and discussion on the manuscript. We would also like to thank all the postdocs that have generously volunteered to share their data.

## Contributions

M.N., R.F.E., P.G., and M.C. conceived and designed the manuscript, curated and interpreted the data. R.F.E. and P.G. acquired the data and M.N. analyzed it. M.N. and M.C. visualized the data. M.C. developed the theoretical framework and supervised the manuscript. M.C. wrote and all authors read the manuscript.

